# The plant circadian clock influences rhizosphere community structure and function

**DOI:** 10.1101/158576

**Authors:** Charley J. Hubbard, Marcus T. Brock, Linda T.A. van Diepen, Loïs Maignien, Brent E. Ewers, Cynthia Weinig

## Abstract

Plants alter chemical and physical properties of soil, and thereby influence rhizosphere microbial community structure. The structure of microbial communities may in turn affect plant performance. Yet, outside of simple systems with pairwise interacting partners, the plant genetic pathways that influence microbial community structure remain largely unknown, as are the performance feedbacks of microbial communities selected by the host plant genotype. We investigated the role of the plant circadian clock in shaping rhizosphere community structure and function. We performed 16S rRNA gene sequencing to characterize rhizosphere bacterial communities of *Arabidopsis thaliana* between day and night time points, and tested for differences in community structure between wild-type (Ws) *vs*. clock mutant (*toc1-21, ztl-30*) genotypes. We then characterized microbial community function, by growing wild-type plants in soils with an overstory history of Ws, *toc1-21 or ztl-30* and measuring plant performance. We observed that rhizosphere community structure varied between day and night time points, and clock misfunction significantly altered rhizosphere communities. Finally, wild-type plants germinated earlier and were larger when inoculated with soils having an overstory history of wild-type in comparison to clock mutant genotypes. Our findings suggest the circadian clock of the plant host influences rhizosphere community structure and function.

## Introduction

In comparison to unvegetated soils, the presence of plants dramatically affects the structure of soil microbial communities. Plant roots affect the physical as well as chemical environment through the exudation of carbon into the rhizosphere zone, which immediately surrounds the roots (Bais et al., 2006; Jones *et al*., 2009; Dennis *et al*., 2010). Rhizosphere microbial community structure is dynamic and changes over the course of plant development (Lundberg *et al*., 2012), in part due to changes in exudation (Chaparro *et al*., 2014). Although much is known about rhizosphere assembly dynamics on longer time scales, there is currently little information regarding assembly dynamics on shorter, diurnal time scales. Further, while plant exudation may “feed-down” and affect microbial community structure, rhizosphere communities can “feed-up” and affect plant performance, by increasing plant access to nutrients (Çakmakçı *et al*., 2001; Chen *et al*., 2002; Richardson *et al*., 2009; Richardson and Simpson, 2011), relieving abiotic stress (Zolla *et al*., 2013), suppressing pathogens (Mendes *et al*., 2011, 2013), altering phenology (Wagner *et al*., 2014; Panke-Buisse *et al*., 2014), and promoting plant growth (Bashan, 1998; Lugtenberg and Kamilova, 2009; Henning *et al*., 2016). Some plant species, such as many legumes, have developmental genetic mechanisms that attract explicitly beneficial nitrogen-fixing rhizobia taxa (Bravo *et al*., 2016). The extent to which plants may attract complex beneficial communities remains largely unclear.

The use of experimental genetic lines available in plant model species may reveal specific genetic paths that affect microbial community structure. Comparing mutant *vs.* wild-type plants of *Arabidopsis thaliana*, Lebeis *et al.* (2015) observed that salicylic acid, an immune signaling molecule, altered rhizosphere bacterial community structure. This finding suggests that genes regulating physiological traits, such as immune response, may play a role in shaping rhizosphere communities. Genes regulating additional physiological traits such as gas-exchange and specifically carbon assimilation may also be worth examining, due to their effects on the photoassimilate pool available for allocation to root growth and for carbon exudation into the rhizosphere. More generally, the comparison of phenotypes between single-locus mutant genotypes with those expressed by wild-type genotypes removes potentially confounding effects of variation segregating elsewhere in the genome, and enables isolation of pathway-specific effects. Naturally occurring large-effect alleles at causal loci could in some cases play a role in shaping microbial community structure in natural plant populations or could be manipulated in crop species to improve plant growth.

Changes in the presence-absence (or abundance) of just a few microbial taxa can affect plant performance, due to the vast number of root-associated microbial cells and functions (Henning *et al.*, 2016). Across diverse ecosystems, community structure and function are related (Tilman *et al.*, 1997; Talbot *et al.*, 2014), including in plant-rhizosphere associations and in cases where microbial community membership changes by one to few taxa. For instance, Henning et al. (2016) observed that the addition of single bacterial taxa to the rhizosphere of *Populus trichocarpa* led to drastic changes in plant growth traits. Similarly, Zolla et al. (2013) observed differences in drought response by *Arabidopsis thaliana* plants grown in soil that differed in community structure as a consequence of overstory history. Thus, differences in rhizosphere community structure can lead to differences in rhizosphere community function as estimated from plant performance.

In the current study, we tested the role of the plant circadian clock in determining rhizosphere community structure and function, where function was measured as plant performance. The circadian clock regulates up to 30% of the transcriptome, and affects diverse processes including patterned fluxes of carbon into (stomatal conductance, carbon assimilation) and out of (exudation) the plant on a diurnal scale (Watt and Evans, 1999; Harmer *et al.*, 2000; McClung, 2006; Covington *et al.*, 2008; Badri and Vivanco, 2009; Harmer, 2009; Greenham and McClung, 2015). We hypothesized that the circadian clock could shape rhizosphere community structure on a diurnal scale, if community structure responds to diurnally patterned fluxes of carbon into the rhizosphere, that is, we anticipated that microbial community structure might vary over the course of 24 hrs. We further hypothesized that rhizosphere communities of plant genotypes harboring clock mutations could differ from wild-type plants, due to differences in physiological phenotypes. Specifically, mutations in the clock genes *TIMING OF CAB EXPRESSION 1* and *ZEITLUPE* lead plants to express altered clock period, or the duration of one circadian cycle (Millar *et al.*, 1995; Kim *et al.*, 2005). As a consequence of altered clock function, clock mutants express distinct physiological phenotypes, including reduced carbon assimilation, chlorophyll content, and stomatal conductance (and thus root water uptake) relative to wild-type plants under 24 hr environmental cycles (Dodd *et al.*, 2004, 2005). Clock misfunction may influence rhizosphere communities, if for instance the reduced flux of carbon into plants influences the flux of carbon exudation (Thornton *et al.*, 2004) or if shifts in plant water use alter soil water potential and nutrient availability and hence the rhizosphere environment (Matimati *et al.*, 2014). Finally, if rhizosphere community structure is altered by mutations in clock genes, then we hypothesize there may be differences in community function in the form of plant performance, in which microbial communities shaped by wild-type genotypes may lead to improved plant performance in comparison to microbial communities found in association with clock mutant genotypes.

## Materials and Methods

### Plant Material and Growth Conditions

To investigate the role of the circadian clock in shaping rhizosphere community structure and function, we used the *Arabidopsis thaliana* accession, Wassilewskija (Ws, CS2360), and two circadian clock period mutants in the Ws background, *TIMING OF CAB EXPRESSION* 1 *(tocl-21)* and *ZEITLUPE (ztl-30). toc1-21* is a short period mutant (∼ 20 hours), while *ztl-30* is a long period mutant (∼28 hours) in free-running conditions (Kevei *et al.*, 2006; Fujiwara *et al.*, 2008). Many prior studies have shown that the resonance between endogenous and environmental cycles affects plant phenotypes and performance (Dodd *et al.*, 2005; Yerushalmi and Green, 2009; de Montaigu *et al.*, 2015; Salmela *et al.*, 2016); the current experiments extend prior research to test effects of the plant host clock on the rhizosphere microbiome.

For each experiment, seeds were surface sterilized using 15% bleach, 0.1% *Tween*, and 84.9% RO H_2_O solution, cold stratified in the dark in 1ml of RO H_2_O for five days at 4°C, and placed in RO H_2_O to germinate in a Percival PGC-9/2 growth chamber (Percival Scientific, Perry, IN, USA) to ensure synchronous germination. Throughout this study, the growth chamber environment was set to 12/12 light-dark cycle (lights came on at 7 A.M. and turned off at 7 P.M.), 22°C/18°C day-night temperature cycles, 40% relative humidity, and photosynthetic photon flux density = 350 μmol photons m^-2^ s^-1^. Upon the observation of root radicles, seedlings were aseptically transferred to 2 inch in diameter pots filled with a mixture of sterilized potting media (N ∼ 400 ppm, P ∼ 90 ppm) and microbial inoculate. To generate our sterilized media, Redi-Earth Potting Mix (Sungro Horticulture, Agawam, MA, USA) was autoclaved twice for 60 minutes. Next, 2ml of microbial inoculate was added to each pot. The microbial inoculate was created by mixing 360ml of RO H_2_O with 40g of soil from the Catsburg region in Durham, North Carolina, USA (36.062294°N, -78.849644°W) and filtered through 1 000μm, 212μm, 45μm sieves, to remove soil nematodes that might negatively impact plant performance (van de Voorde *et al.*, 2012). The Catsburg region has a well-documented history of *A. thaliana* occurrence, which has been naturalized in this region (Mauricio, 1998). Soil from the Catsburg region and our sterilized potting mix was characterized at the Colorado State Soil-Water-Plant Testing Lab (Fort Collins, CO, USA); of greatest relevance to microbial growth, the Catsburg and potting soils had similar pH values (5.4 *vs.* 5.3, respectively). Following germination, seedlings were thinned to one plant per pot, and pots were watered at 7 A.M. daily.

### Experimental Design

#### Experiment 1: Temporal Changes in Rhizosphere Community Structure

To determine if rhizosphere bacterial communities are diurnally dynamic, replicates of wild-type Ws plants were grown for four weeks as described above. Starting at 6 A.M. on July 21 and ending at 6 A.M. on July 22, 10 replicates were selected at random and harvested every 6 hrs for rhizosphere soil by separating the roots from the rosette (N = 50), removing closely adhering soil particles from the roots as described in Bulgarelli et al. (2012), and storing the samples at -80°C.

#### Experiment 2: Candidate Drivers of Rhizosphere Community Structure

To characterize the effects of circadian period misfunction on rhizosphere bacterial community structure, 10 replicates of Ws, *toc1-21*, and *ztl-30* genotypes were planted in a fully randomized design and grown for four weeks. Rhizosphere samples were collected as described above at 6 P.M. on July 21 and stored at -80°C. All samples were collected prior to visible signs of bolting, or the transition from a vegetative to a reproductive state, to avoid confounding effects of plant developmental stage (Lundberg *et al.*, 2012; Chaparro *et al.*, 2014). At the end of this experiment, we collected additional rhizosphere soil from 4 replicates of each of the three genotypes to generate the inoculum for *Experiment 3.*

#### Experiment 3: Rhizosphere Community Feedbacks on Plant Performance

To test if rhizosphere microbiomes assembled by the three plant genotypes had differential effects on plant performance, we synchronously germinated seeds of the Ws genotype and planted these seeds in sterilized soil media inoculated with soil slurry generated by the Ws, *toc1-21*, or *ztl-30* genotypes and collected at the end of *Experiment 2* (N = 60; 20 replicates × 3 inoculates). To determine the effects of the rhizosphere microbiome treatment on plant performance, rosette diameter was measured weekly for three weeks. In a second experiment, we allowed seeds to germinate naturally in sterilized soil media inoculated with the same soil slurries *(N* = 60; 20 replicates x 3 inoculates). For this experiment, seeds were checked daily for germination as estimated from the first observation of cotyledons.

### DNA Extraction and Amplicon Sequencing

To extract microbial DNA, rhizosphere samples were placed into 15ml Nunc Conical Centrifuge Tubes (Thermo Scientific, Waltham, MA, USA) containing 3ml of phosphate-buffered saline (PBS), and then agitated for 15 minutes to separate soil particles from plant roots as described in Bulgarelli *et al.* 2012. Plant roots were then removed with sterilized forceps and the samples were centrifuged for 15 minutes at 3000 rcf. The supernatant was discarded, and 0.25g of the pellet was put into bead tubes from the Mobio Power Soil DNA Isolation Kit (Mobio Laboratories, Carlsbad, CA, USA) using sterilized disposable spatulas. DNA was extracted from each sample following the manufacturer’s instructions. With each round of extractions, a soilless blank was included as a negative control. At the end of each round, PCR was performed to ensure sufficient DNA yields and reagent sterility.

DNA extracts were sent to the Marine Biological Laboratories (Woods Hole, MA, USA) for amplicon library preparation of the V4V5 region of the 16S rRNA gene using the 518F and 926R primers (Huse *et al.*, 2014). Sequencing was performed on the Illumina MiSeq platform (Illumina, San Diego, CA, USA) as described in Nelson et al. (2014). Sequence reads were demultiplexed and quality filtered (Phred score ≥ 20, chimera removal by ChimeraSlayer) using QIIME 1.9.1, uclust was used to perform open reference OTU picking at 97% sequence similarity using the Greengenes database (ver. 13.8), and all singletons were removed to avoid the possibility that a sequencing error was called as an OTU (Caporaso *et al.*, 2010; Bokulich *et al.*, 2012; McDonald *et al.*, 2012; Edgar, 2010; Haas *et al.*, 2011). We rarefied to 100 000 reads per sample to ensure common sampling effort. All sequences have been deposited into the Short Read Archive (SRA) under accession number SRA579608.

### Sequencing data analyses

To describe rhizosphere community structure, we generated Jaccard (presence-absence analysis) and Bray-Curtis (abundance analysis) dissimilarity matrices, and Shannon diversity estimates in QIIME (Caporaso *et al.*, 2010). For *Experiment 1*, we used adonis to determine if rhizosphere community structure differed between day (6 P.M.) and night (6 A.M.) time points. To test if shifts in community structure were consistent between day and night time points, we used Pearson’s correlation coefficients to compare the percent change in OTU abundance between 6 A.M. and 6 P.M. on July 21 and the percent change in OTU abundance between 6 P.M. on July 21 and 6 A.M. on July 22, for OTUs with more than 100 reads per sample to avoid potentially confounding effects of low-abundance taxa. For *Experiments 2 & 3*, we used one-way ANOVAs and Tukey’s HSD *post hoc* comparison tests using the *car* and *agricolae R* packages to characterize differences in Principal Coordinates between genotype along axis 1 and to determine differences in plant performance between soil treatments (Fox *et al.;* de Mendiburu, 2016). Moreover, OTUs were split into common (> 500 reads) or rare (< 500 reads) categories, and presence-absence analyses and abundance analyses were performed again on the split datasets to determine if effects of clock genotype were detected using common or rare microbial taxa alone. Further, sequence data was reanalyzed without rarefaction using the R package *Phyloseq*, to determine if results were consistent in the absence of rarefaction (McMurdie *et al.,* 2014). Results were similar regardless of rarefaction, that is, the effect of host plant genotype was significant for both binary Jaccard (p = 0.001) and Bray-Curtis dissimilarity (p = 0.001) analyses with and without rarefaction; here, we present the results of analyses based on rarefaction. All plots were generated using the R package *ggplot2* (Wickham, 2009).

To identify OTUs that explain observed differences in plant performance due to soil overstory history in *Experiment 3*, we used the *indicspecies* package for indicator value analysis in R 3.0.3 and *LefSe* on the galaxy web platform (Cáceres and Legendre, 2009; Dufre□ne and Legendre, 1997; Segata *et al.*, 2011; R Core Team, 2013). Indicator value analysis (IVA) has been used commonly in ecological studies to ascertain species that underlie treatment or site differences (Dufre_ne and Legendre 1997), and is used here to test which OTU(s) is(are) specific to a given level of a factor (e.g., present / abundant in the rhizospheres of Ws replicates and absent from *toc1-21* and *ztl-30* rhizospheres). Notably, the calculation of IVA weights presence-absence and abundance, and as such may be sensitive to rare taxa. *LefSe* performs linear discriminant analysis on sequence data to identify marker taxa that underlie treatment differences, and is weighted preferentially by abundance differences of more common taxa. Because rare OTUs contributed to microbiome differences between host plant genotypes, we used both IVA and *Lefse.* Finally, we coarsely estimated microbial community size by dividing the quantity of extracted DNA using a Qubit (ThermoFisher Scientific Waltam, MA, USA) by the mass of soil used for each extraction to determine if microbial community size influenced plant performance in *experiment 3.*

## Results

### Sequencing Results

For *Experiment 1*, after quality filtering, chimera removal, OTU picking, outlier sample filtering and rarefaction to 100 000 reads per sample (**Supplemental Figure 1a**), there was a total of 3 700 000 high quality reads out of 10 250 881 raw reads. For *Experiment 2*, after similar processing, but rarefaction to 116 000 reads per sample (**Supplemental Figure 1b**) there was a total 2 668 000 high quality reads out of 6 487 790 raw reads. The number of reads after each processing step can be found in **Supplemental Tables 1** and **2.**

### Experiment 1: Temporal Changes in Rhizosphere Community Structure

We found significant differences in rhizosphere community structure between the communities collected at 6 P.M. (day) and both of the two 6 A.M. (night) collections (P = 0.001 contrasting 6 P.M. with the first collection at 6 A.M. on July 21 or P = 0.008 contrasting 6 P.M. with the collection at 6 A.M. on July 22; Figure 1a). The percent change in OTU abundance between 6 A.M. on July 21 and 6 P.M. was positively correlated with the percent change in OTU abundance between the 6 A.M. on July 22 and 6 P.M. time points (r = 0.423, P < 0.001). This relationship suggests that the abundance of many common OTUs shift in a similar manner between day and night time points (Figure 1b).

**Figure 1:**
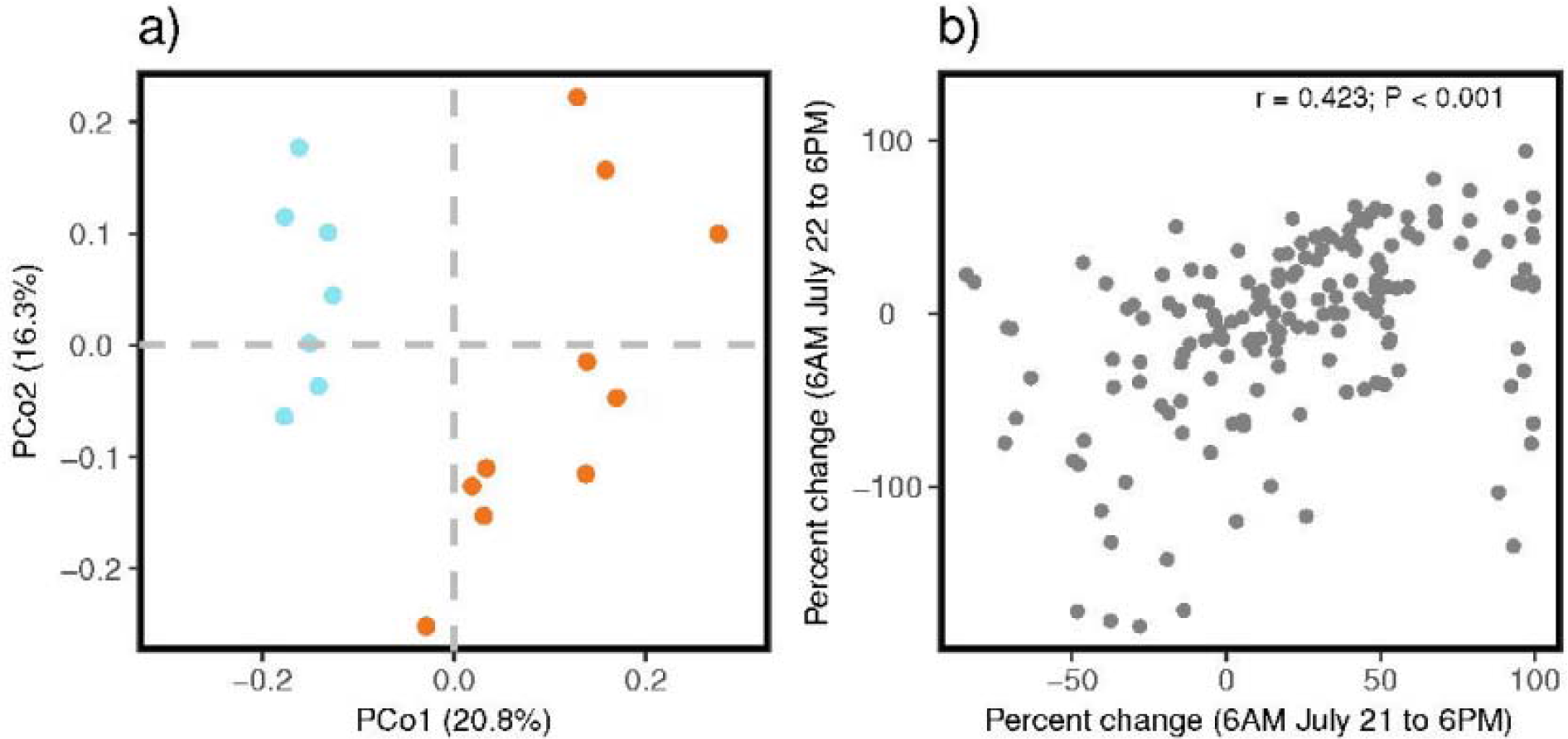
Day *vs.* night influences rhizosphere community structure. **a)** Principal Coordinate Analysis of Bray Curtis dissimilarities *(n* = 17). Differences between the 6 P.M. (day, orange) and 6 A.M. (night, blue) time points were significant at P = 0.001. **b)** Bivariate relationship between percent change in OTU abundance from 6 A.M. to 6 P.M. July22 and percent change in OTU abundance from 6 A.M. on July 21 to 6 P.M. (r = 0.423; P < 0.001).

### Experiment 2: Candidate Drivers of Rhizosphere Community Structure

Rhizosphere community composition (presence *vs.* absence of taxa), abundance, and diversity differed among genotypes (Figure 2, **Supplemental Figure 2**). From Jaccard presence-absence analysis, Ws, *toc1-21*, and *ztl-30* rhizosphere communities were significantly different from one another (P < 0.001; Figure 2a). PCo1 describes the effect of progressive clock changes between short (lowest PCo values) vs. wild-type (intermediate values) vs. long (highest values) endogenous period lengths of the plants on rhizosphere community composition. Bray-Curtis relative abundance analysis revealed differences between the rhizosphere communities of the three clock genotypes, where *toc1-21* communities were different from both Ws (P = 0.02) and *ztl-30* (P = 0.04) communities, while Ws and *ztl-30* communities were not significantly different from one another (P = 0.39; Figure 2b). This result suggests that a period length shorter than 24 hrs specifically alters abundances of OTUs within the rhizosphere community. Likewise, the *toc1-21* rhizosphere communities showed significantly reduced richness and evenness based on Shannon’s diversity index in comparison to Ws (P = 0.03; Figure 2c).

**Figure 2:**
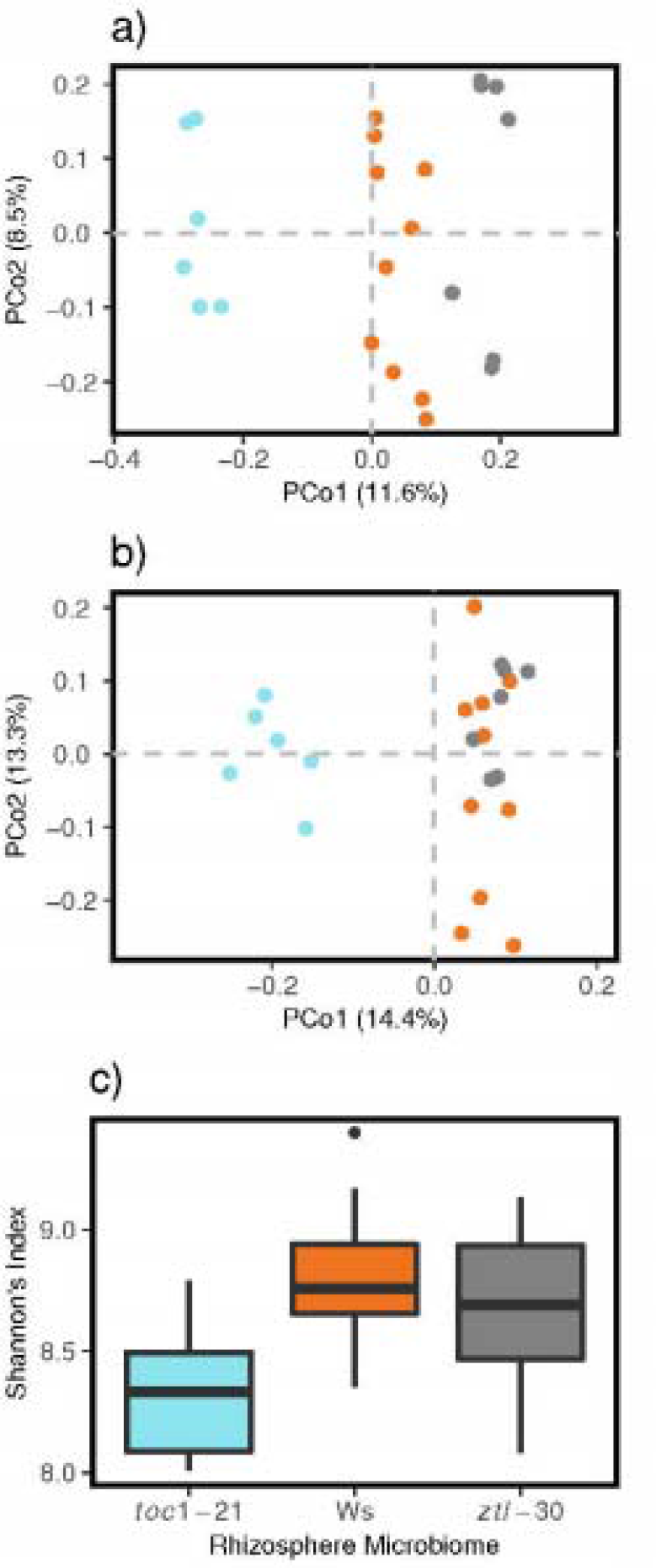
Clock function in *A. thaliana* alters rhizosphere community composition, abundances, and diversity. (**a**) Principal coordinate analysis of Jaccard dissimilarities, where rhizosphere communities of Ws are represented by orange circles, *toc1-21:* blue circles, and *ztl-30:* gray circles (n = 23). Rhizosphere community composition differs significantly between clock genotypes (P = 0.001). (**b**) Principal coordinate analysis of Bray-Curtis dissimilarities (n = 23). OTU abundances differ significantly between *toc1-21* and both the Ws and *ztl-30* genotypes (P = 0.001). (**c**) Mean Shannon diversity index. The top and bottom of boxes represent the 75^th^ and 25^th^ percentiles, respectively. Whiskers represent 1.5 times the interquartile range. One-way ANOVA and Tukey’s *post hoc* comparisons indicate significant differences between *toc1-21* and both the Ws and *ztl-30* genotypes (P = 0.03).

To clarify the contributions of rare vs. common OTUs to host plant genotype differences, we analyzed the data when culled to different minimum read numbers. When culling to a minimum read number of 1 000 for an OTU (or approximately 1% of the community), Jaccard and Bray Curtis dissimilarities were significant, indicating that common taxa contribute at least partially to observed differences among the three host plant genotypes in the presence-absence of taxa (Figure 3a) and to differences in OTU abundance between *toc1-21* and both Ws and *ztl-30* (Figure 3b). Communities culled to OTUs with between 1-500 reads showed significant differences in both composition and abundance, indicating that rare microbial taxa respond to plant genotype (Figure 3cd). In particular, when data for rare OTUs are analyzed, the distinction between Ws vs. *ztl-30* becomes significant (p = 0.001) (cf Figure 2b vs. Figure 3d).

**Figure 3:**
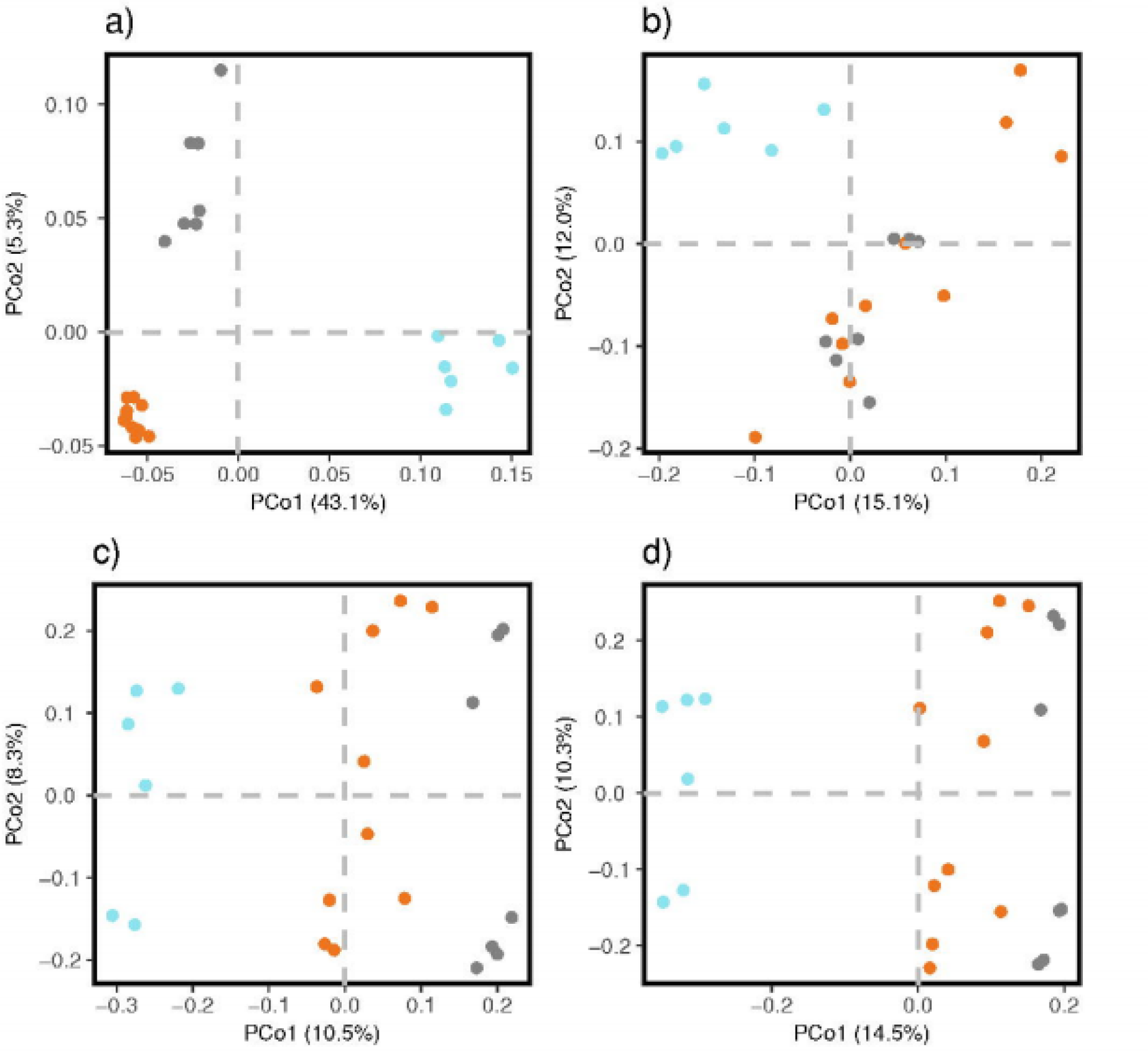
Plant genotype influences both common (> 500 reads) and rare (1-500 reads) rhizosphere taxa, but differences between the Ws and *ztl-30* rhizosphere communities are attributable to rare taxa. (**a**) Principal coordinate analysis of Jaccard dissimilarities of common taxa, where rhizosphere communities of Ws are represented by orange circles, *toc1-21:* blue circles, and *ztl-30:* gray circles *(n* = 23). Rhizosphere community composition differs significantly between clock genotypes (P = 0.001). (**b**) Principal coordinate analysis of Bray-Curtis dissimilarities of common taxa (n = 23). OTU abundances differ significantly between *toc1-21* and both the Ws and *ztl-30* genotypes (P = 0.001). (**c**) Principal coordinate analysis of Jaccard dissimilarities of rare taxa (n = 23). Rhizosphere community composition differs significantly between clock genotypes (P = 0.001). (**d**) Principal coordinate analysis of Bray-Curtis dissimilarities of rare taxa (n = 23). OTU abundances differ significantly between Ws and both *toc1-21* and *ztl-30* genotypes (P = 0.001).

Combined, the IVA and *Lefse* analyses identified a total of 13 indicator OTUs associated with the Ws rhizosphere (Figure 4), 12 indicator OTUs associated with the *toc1-21* rhizosphere (**Supplemental Tables 7, 9**), and 12 indicator OTUs associated with the *ztl-30* rhizosphere (**Supplemental Table 8, 9**). Because IVA is more sensitive to rare taxa, the two methods select somewhat different OTUs as biomarkers of host plant genotype. Notably however, there is significant taxonomic overlap between the OTUs identified by IVA and *Lefse.* That is, taxa identified by IVA are phylogenetically related to those identified by *Lefse*, or vice versa. For instance, of the 13 indicator OTUs associated with the Ws rhizosphere, six taxa were members of the phylum Acidobacteria (IVA: DS-100;o_;f_;g_, llb;f_;g_, PAUC26;f_;g_; *Lefse:* Acidobacteria, Solibacterales;f_;g_, iii1_15;f_;g_) and two taxa were members of the Chloroflexi (Indicator species analysis: Anaerolineae;o_;f_;g_, *Lefse:* Chloroflexi). From previous studies, both phyla and two taxa, *Agromyces* and *Cellulomonas*, have been previously described as growth promoting (Egamberdiyeva and Höflich, 2002; Kuffner *et al.*, 2008; Chen *et al.*, 2014; Kielak *et al.*, 2016). Finally, community size as estimated from DNA per unit soil mass did not significantly differ across clock genotypes (P = 0.11).

**Figure 4:**
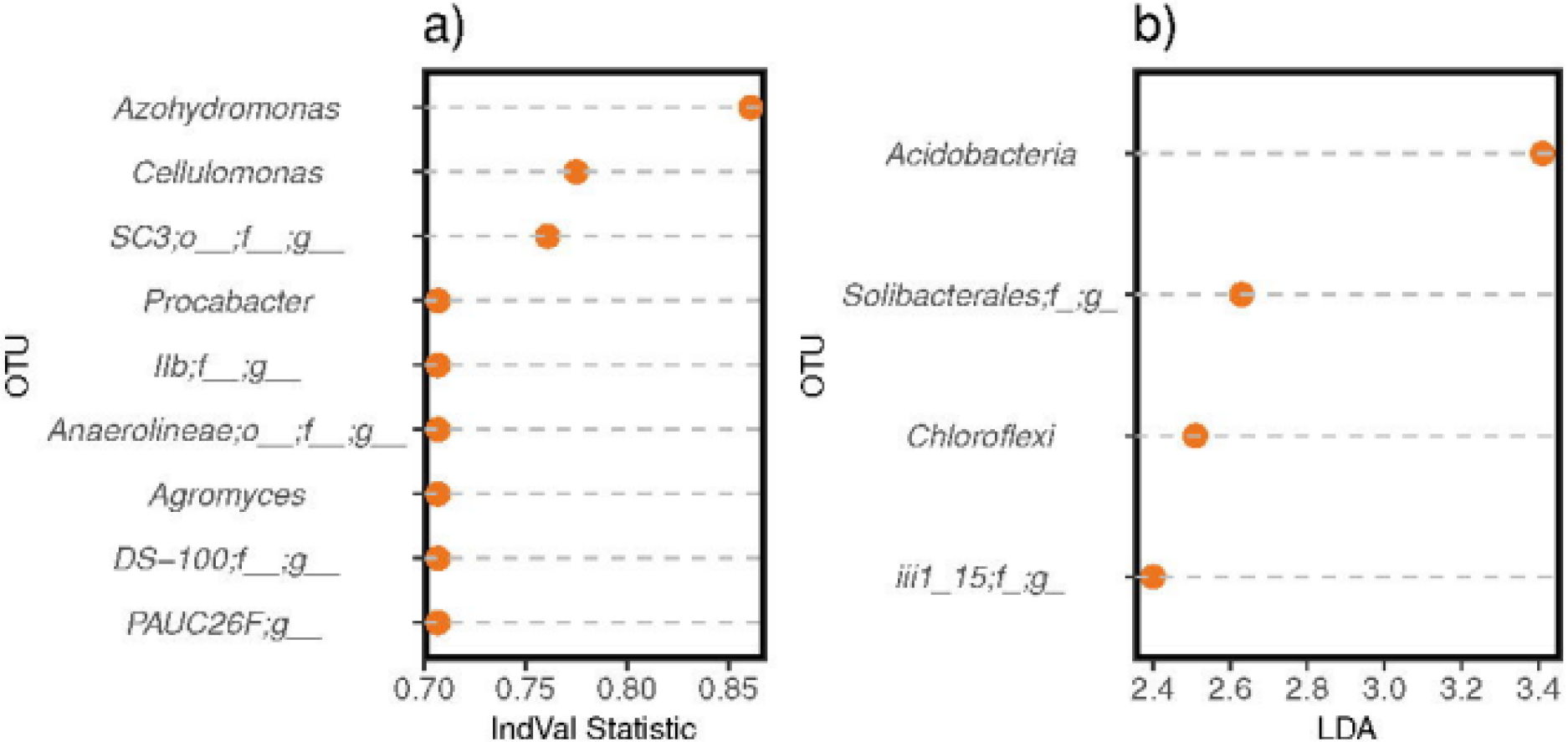
Indicator taxa based on (**a**) Indicator value analysis and **(b)** *Lefse.*

### Experiment 3: Rhizosphere Community Feedbacks on Plant Performance

Soil overstory history had a significant influence on early plant performance (Figure 5). Wild-type plants grown in a soil with a history of Ws plants had significantly larger rosette diameters than plants grown in soils with a history of *toc1-21* and *ztl-30* after 1 week (19.4% and 14.4%, respectively; P = 0.002) and 2 weeks of growth (10.8% and 8.3%, respectively; P = 0.04). However, at the end of three weeks of growth, Ws plants grown in soils conditioned by each of the clock genotypes were only marginally different in size (P = 0.11). In a germination experiment of similar design (in which seedlings were not transplanted but instead germinated directly on soil), Ws seeds in pots with Ws inoculum germinated an average of 5.2 days earlier than seeds planted into pots with *toc1-21* (P = 0.002) inoculum and 5.7 days earlier than those planted into *ztl-30* (P = 0.024) inoculated pots (P < 0.001; Figure 5b).

**Figure 5:**
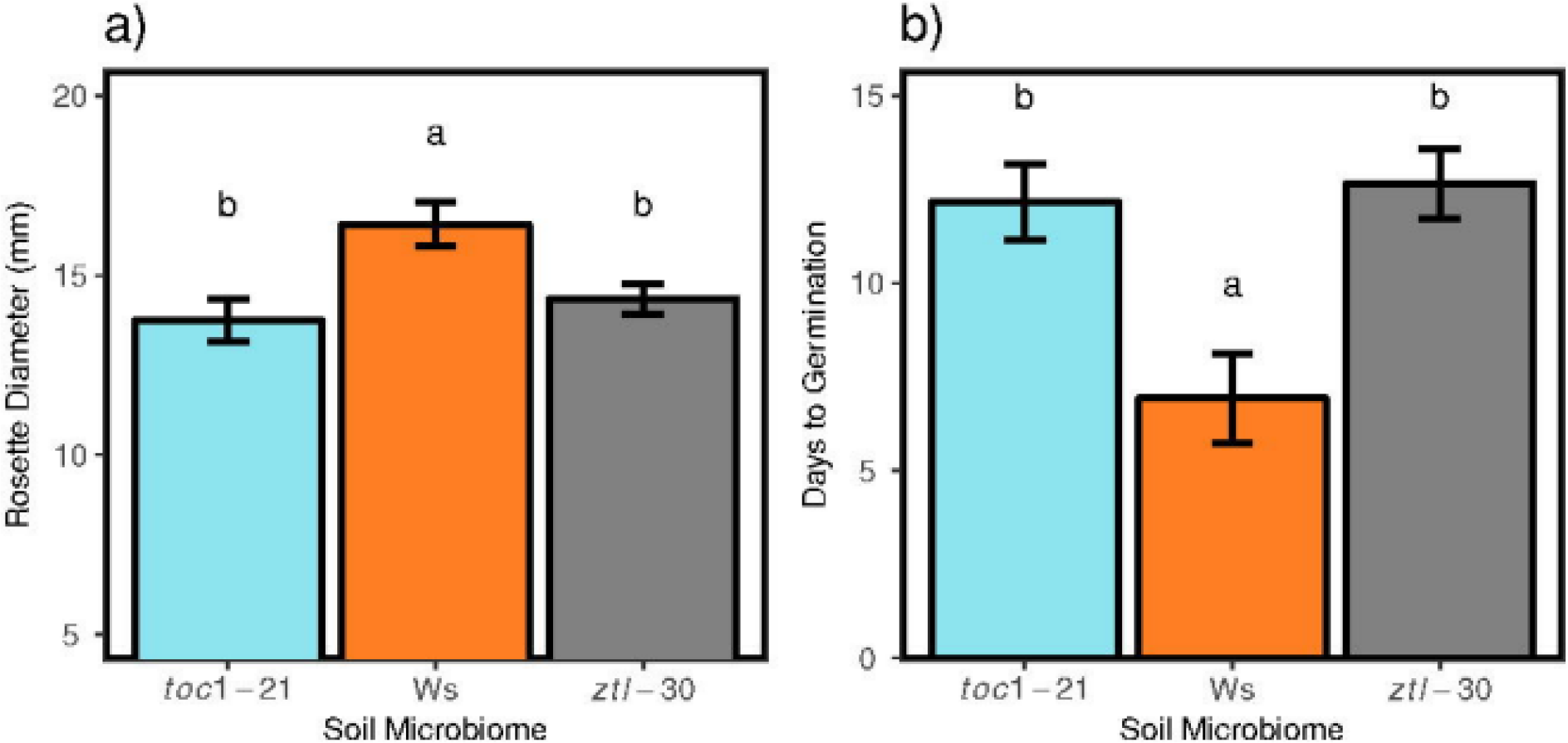
Effects of soil overstory history on plant growth. Letters denote significant differences between soil treatments based on Tukey’s Honest Significant Differences *post hoc* comparisons. (**a**) At Week 1 (n = 60), plants grown in a soil with a history of Ws had significantly larger rosette diameters than plants grown in soils with a history of *toc1-21* or *ztl-30* (P = 0.002). (**b**) In a separate experiment where seeds were not germinated synchronously (n = 35), seeds sowed onto a soil with a history of Ws occurrence germinated significantly earlier than seeds sown into soils with a history of *toc1-21* or *ztl-30* genotypes (P < 0.001).

## Discussion

The rhizosphere microbiome has been referred to as the “second genome” of plants or the extended phenome (Berendsen *et al.*, 2012). In part, these names reflect the role of the rhizosphere microbiome in determining plant performance. Empirical studies suggest complex feedbacks between plants and microbes, under which plant species may modulate rhizosphere community structure via carbon exudation and under which microbes may alter plant phenotypes directly or via ecosystem services such as nutrient accessibility (Bulgarelli *et al.*, 2013). The mechanisms by which different plant genotypes may influence rhizosphere community structure remain largely unclear, as are the effects on plant performance of rhizosphere microbiomes selected by the plant host genotype (Heath and Tiffin, 2007; Panke-Buisse *et al.*, 2014; Lebeis *et al.*, 2015). Understanding plant-rhizosphere microbiome interactions is agroecologically relevant because rhizosphere communities can strongly influence plant fitness and biomass, which can in turn inform evolutionary studies of adaptation, conservation, and agronomic practices (Pérez-Jaramillo *et al.*, 2016). In this study, we tested the role of the plant circadian clock as a mediator of plant-rhizosphere microbiome interactions. We hypothesized that 1) rhizosphere community structure may be temporally dynamic, if rhizosphere taxa may respond to diurnally patterned fluxes of carbon, water, or nutrient availability into the rhizosphere (or other diurnally patterned plant phenotypes). 2) We further hypothesized that clock misfunction would play a role in shaping community structure, because differences in plant physiology attributable to genotype would lead to differences in rhizosphere community structure. 3) Finally, we hypothesized that differences in rhizosphere community structure attributable to plant genotype could lead to differences in community function with regards to plant performance.

The composition of plant-associated microbiomes is known to shift on long time scales, such as across seasons or across developmental stages of the plant host (Lundberg *et al.*, 2012; Chaparro *et al.*, 2014; Wagner *et al.*, 2016). The short duration of many microbial life cycles means that microbial community composition may also respond to more rapid changes in the environment. Yet, it remains unclear if the community composition of microbes found in association with plants changes on short timeframes, such as across day-night transitions. We observed diurnally patterned shifts in rhizosphere community structure. For instance, we observed shifts in community structure between day (6 P.M.) and night (6 A.M.) time points, and we observed that these shifts in community structure were consistent between the day and both night time points (Figure 1). Difference in community structure observed between the two time points may reflect the effects of day *vs.* night conditions and carbon, water, or nutrient availability in the rhizosphere. Several prior studies have shown that the concentration of certain exudates varies over the course of day (Badri and Vivanco, 2009; Watt and Evans, 1999). For instance, Iijima *et al.* (2003) observed higher rates of mucilage exudation at night, while other studies have observed higher prevalence of flavonoids and catechin during day conditions (Hughes *et al.*, 1999; Iijima *et al.*, 2003; Tharayil and Triebwasser, 2010). Further, rhizosphere water is depleted diurnally, depending on root and soil hydraulics (Sperry *et al.*, 1998), and the transpiration stream increases nutrient flow (Matimati *et al.*, 2014), potentially depleting soil nutrients in the rhizosphere zone. Therefore, rhizosphere taxa and populations may vary in abundance depending on soil resource availability, leading to our observed differences in community structure between day and night time points. Future experiments should be designed to tease apart the relative influence of root exudates, water dynamics, and nutrient uptake within the rhizosphere on microbes.

Specific plant genes play important roles in shaping rhizosphere community structure (Bravo *et al.*, 2016), and here we observed that circadian clock genes significantly influence rhizosphere community structure. In the current study, plant genotype explained 19.1% of the variation in community composition (presence *vs.* absence of taxa), 21.7% of the variation in community relative abundances, and brought about differences in community diversity between short *(toc1-21) vs.* longer (Ws, *ztl-30)* period genotypes (Figure 2). These differences in community structure explained by clock genotype surpass variation explained by genotype in previous studies of the influence of plant genotype on rhizosphere community structure (Lundberg *et al.*, 2012; Bulgarelli *et al.*, 2012; Peiffer *et al.*, 2013; Lebeis *et al.*, 2015). Our results thus suggest that circadian clock misfunction has a strong influence on rhizosphere community structure. The large percent variance explained may arise from the pervasive transcriptomic and phenotypic effects of clock misfunction on the plant host, or potentially the microbial inoculant used here is one that amplifies the effect of host genotype, as demonstrated in other studies (Weinert *et al.*, 2011; Peiffer *et al.*, 2013).

*TIMING OF CAB EXPRESSION 1* had a particularly pronounced impact on rhizosphere community structure, because strong mutant alleles in this gene led to changes in community composition, abundance, and diversity (Figure 2). On the other hand, disruption of *ZEITLUPE* had less of an influence on shaping rhizosphere community structure, with its effect limited to differences in OTU presence-absence relative to Ws (Figure 2). One possible explanation for the asymmetric effects of clock misfunction is that long period lengths theoretically enable better phase adjustment to dawn, such that period lengths shorter than 24 hrs may have more detrimental fitness consequences in nature (or in this case lead to greater deviations in rhizosphere microbial community structure) in comparison to period lengths greater than 24 hrs (Johnson and Kondo, 1992; McClung, 2006; Kevei *et al.*, 2006; Hotta *et al.*, 2007). Regardless of the exact mechanisms, clock misfunction and the mismatch between endogenous plant cycles and exogenous cycles affected aspects of microbial community structure.

As in any ecosystem, there is a link between community structure and function (Tilman *et al.*, 1997). Several studies have illustrated this relationship in plant-rhizosphere microbiome interactions, where differences in plant performance can be attributed to differences in rhizosphere community structure (Mendes *et al.*, 2011; Zolla *et al.*, 2013; Mendes *et al.*, 2013; Wagner *et al.*, 2014). Here, differences in community structure brought about by mutations in circadian clock genes led to differences in plant performance among wild-type Ws plants grown in soils with differing plant genotype overstory histories (Figure 5). Ws plants performed best when exposed to an inoculum from soils in which wild-type rather than clock mutant genotypes had been grown. We observed differences in the timing of germination, where Ws seeds sown in soils with a history of Ws occurrence germinated earlier. In comparison to untreated soil, autoclaved soil, such as that used here, differs in both chemical and physical properties and reflects a novel and possibly more stressful environment for plants (Trevors, 1996; Brulé *et al.,* 2001; Lau and Lennon, 2011); differences in germination observed here may therefore reflect that the microbes from Ws-conditioned soil enable normal germination under the novel autoclaved soil conditions (rather than an acceleration of germination timing under natural conditions *per se)* (Lau and Lennon, 2011; Mahmood *et al.*, 2014). Beyond germination timing, we observed that wild-type plants were also larger when grown in soils with a history of Ws rather than mutant genotype growth. These findings from two experiments in which soils were independently conditioned by Ws vs. mutant genotypes suggest first that plants can select explicitly beneficial soil communities that improve initial offspring performance, and second that disruption of these communities by mutations in clock genes adversely affects initial offspring phenology and growth.

While additional research is required to ascertain causality, community composition patterns and indicator analyses provide hypotheses as to which OTUs may lead to these differences in performance (Cáceres and Legendre, 2009; DeAngelis *et al.*, 2015). Here, we identified 13 indicator OTUs associated with Ws rhizosphere. Parallel to the differences in microbial community structure among host plant genotypes (Figures 2 and 3), one possibility is that rare OTUs underlie differences in plant performance observed between the Ws and *ztl-30* rhizosphere microbiomes, while rare and common OTUs could contribute to plant performance differences observed between the Ws and *toc1-21* microbiomes. Rare OTUs could affect plant performance via so-called indirect effects, such as facilitation of or competition with explicitly plant growth-promoting microbes, while common microbes could promote plant growth through direct interactions (Saleem *et al.*, 2016). Specifically in regard to the indicator species analyses (IVA and *Lefse)*, the presence and abundance of Acidobacteria (Kielak *et al.*, 2016), Chloroflexi (Chen *et al.*, 2014), *Cellulomonas* (Egamberdiyeva and Höflich, 2002), and *Agromyces* (Kuffner *et al.*, 2008) in the Ws rhizosphere may explain the differences in plant size between rhizosphere treatments, as these OTUs have been previously associated with plant growth promotion.

In sum, we have shown that the plant circadian clock shapes rhizosphere community structure, particularly the presence of rare taxa. Further, this plant genetic driver of community assembly also influences community function, as estimated from plant performance. Because community structure may shift in response to day and night conditions, future characterizations of the rhizosphere should account for differences in community structure due to the timing of rhizosphere collection. Moreover, more work needs to be done investigating the role of other pertinent clock genes and loci that regulate plant physiology, as these may also shape rhizosphere community structure and function.

## Acknowledgements

This work was supported by the National Science Foundation grants (IOS-1444571) to C.W., B.E.E., L.M., and a Wyoming INBRE sequencing and bioinformatics award to C.H. and C.W. We thank the Powells’ for allowing us to collect soil from their property and Lindsay Leverett for collecting soil from the Catsburg site.

## Conflict of Interest

The authors declare no conflicts of interest.

